# Feedback processing in attack and defense conflicts: a neurophysiological study

**DOI:** 10.1101/2021.06.25.449933

**Authors:** Tiago O. Paiva, Rui C. Coelho, Rita Pasion, Eva Dias-Oliveira, Carsten K. W. De Dreu, Fernando Barbosa

## Abstract

Despite being extensively modelled to explore decision making processes in economic tasks, there are no studies exploring the neurophysiological correlates of the Predator-Prey Game, a game theory paradigm designed to operationalize attack and defence conflicts. In the present study, we explored the relationship between the ERP components associated with feedback, namely feedback-related negativity (FRN) and feedback -elated P3b, and investment in an adapted version of the Predator-Prey Game (PPG), operationalizing attack and defence. Forty-seven (28 female) community-dwelling volunteers without history of neurological disease were recruited. Participants played the PPG game both as attackers and defenders while EEG signal was recorded with a 128 channels sensor net. Behavioural results showed that individuals tend to invest more and decide faster when playing in defence, rather than in attack. Electrophysiological data showed FRN to be sensitive to the valence of feedback, with increased amplitudes of FRN loss feedbacks. On the other hand, the P3b amplitudes were sensitive to the role, with increased amplitude for attack when compared with defence. The integration of the behavioural and ERP findings supports the theoretical model positing that attack elicits more deliberate and less automatic brain processes than defence.

## 1. Introduction

Although social conflict poses several consequences for human life, studies on the brain mechanisms underlying conflict processing are scarce. Broadly speaking, social conflicts can result from incompatibilities between two (or more) individuals or groups over several aspects of life such as resources, expectations, values, motivations or goals (Koban et al., 2010). Stemming from the observation that conflicts between nonhuman animals follow a structure of attack and defense (e.g., Boehm, 2012; Wrangham, 2018), de Dreu and Gross (2019) suggest that most of human interpersonal conflict follow a similar structure where one side is perceived as a ‘threatening aggressor’, who’s actions trigger some form of aggressive defense (Chambers et al., 2006).

The multidisciplinary study of interpersonal conflict often relies in behavioral game theory to model decisions, strategies, and outcomes in attack and defense (e.g., De Dreu et al., 2019). The operationalization of game theory is performed in the form of games that define a set of strategies for two or more players and allow for the mathematical assessment of the expected utility of the combination of each possible strategy within the game. These games often follow a symmetric structure for conflict, meaning that switching positions between players do not change the preferred choice strategy nor the motivation underlying that strategy (De Dreu & Gross, 2019). The Prisoner’s Dilemma is a classic example of a symmetric game theory formulation of conflict. In this game each of two players, A and B, must decide between two possible actions, cooperate or defect. There are four possible outcomes depending on the combination of the actions selected by the two players: (1) mutual cooperation, where both players cooperate; (2) mutual defection where both players defect; (3 and 4) where one of the players decides to cooperate and the other defects. The best outcome is achieved when one’s decides to defect while the other player decides to cooperate and the worst whenever the player decides to cooperate while the other defects. In between, mutual cooperation is more beneficial to both players than mutual defection. In this formulation defection has the potential to both maximize personal game while protecting against possible exploitation for both players (Coombs, 1973; De Dreu & Gross, 2019).

However, operationalizing conflict in the form of attack and defense follows an asymmetric structure where distinct players have distinct motivations. The Predator-Prey context (De Dreu et al., 2015) is an example of asymmetrical game theory formulation of attack and defense. As described in de Dreu and Gross (2019), in this game the attacker must decide how much to invest in attack (*x*) from a given initial endowment *e* (with 0 ≤ *x* ≤ *e*) while the defender must decide, at the same time, how much to invest in defense (*y*) out of the initial endowment *e* (with 0 ≤ *y* ≤ *e*). If the attacker invests more than the defender (*x > y*) the attacker gains the remaining of both players initial endowment [(*e-x*) + (*e-y*)] while the defender is left with nothing. If the attacker fails to invest more that the defender (*x* ≤ *y*) than both players gain the remaining amount from the initial endowment, that is the attacker gets *e-x* and the defender *e-y*. In this formulation, while attackers invest to maximize reward, defenders invest to protect against exploitation and a change in role is associated with a change in both the motivation to invest and corresponding strategy.

The psychological determinants of attack and defense conflicts can be conceptualized in terms of the behavioral approach-avoidance theory (De Dreu & Gross, 2019). In this formulation, attack is linked with the behavioral activation system modulated by the mesolimbic dopaminergic system, where positive emotions trigger reward seeking behaviors. Defense on the other hand is linked with the behavioral inhibition system modulated by the serotoninergic pathway, where negative emotion trigger behavioral response to threatening cues aimed at avoiding potential loss (De Dreu & Gross, 2019; Gray, 1990). Attack and defense may also differ in the recruitment of a brain network responsible for top-down control, including the inferior frontal gyrus, the dorsolateral and orbitofrontal gyrus, and the anterior cingulate gyrus (Braver, 2012; De Dreu & Gross, 2019; Dosenbach et al., 2008). The greater activation of prefrontal control brain regions during attack (e.g., De Dreu et al., 2015) lends support for the notion that attacking recruits more top-down control than defence, and therefore is more deliberate, and less spontaneous than defence (De Dreu & Gross, 2019).

Two factors may explain the increased automaticity of defence when compared with attack. The first one stems from the Predation Theory (e.g., Taylor, 1984), that attributes increased biological relevance to defence against predation when compared with predation, thus favouring the attainment of automatic and this faster defence responses. The second has to do with the strategic computations necessary to achieve success in the Predator-Prey contest. While attackers maximize gains by mismatching the defender’s level of competitiveness, defenders minimize loss by matching the attacker’s level of competitiveness. Given that mismatching responses is cognitively more demanding than matching responses, attack may putatively elicit increased cognitive control and deliberation (De Dreu & Gross, 2019).

The distinct strategies underling attack and defence may also lead to differential feedback processing when playing as “attacker” or as “defender”. Feedback can be conceptualized as sensitive information reflecting the outcome of an action (Luft, 2014). Either by visual input on the outcome of a specific action (e.g., the number of pins knocked by a bowling ball) or by verbal information (e.g., the bowling score display on the board) feedback is fundamental for both performance monitoring and improvement (Masters et al., 2009), and plays a crucial role in strategy learning in interpersonal interactions. As an example, feedback manipulation has been effective in modulating the level of cooperation in the iterated prisoner’s dilemma (e.g., Monterosso et al., 2002) and fairness orientation in the iterated Ultimatum Game (Bieleke et al., 2017). Given that attack and defence are asymmetrically modelled in the Predator-Prey contest, it is plausible that feedback is processed differently by “attackers” and “defenders”.

The Event-Related Potentials (ERP) technique has been extensively used to assess the brain mechanisms underlying feedback processing (e.g., Chen et al., 2017; Constantino et al., 2021; Luft, 2014). An ERP reflects brain electrical responses synchronized with a specific stimulation and/or event (e.g., feedback from a specific action). The typical ERP waveform represents a composite of distinct components that are modulated by distinct perceptive, emotional, and attentional brain mechanisms within the information processing stream (Luck, 2014). In the context of feedback, two main ERP components have contributed to the understanding of the brain mechanisms underlying feedback processing: (a) the Feedback Related Negativity (FRN) and (b) the P3b (e.g., Chen et al., 2017; Constantino et al., 2021; Luft, 2014). The FRN component is a negative deflection captured in the frontocentral electrodes in between 200 to 350 ms post feedback presentation (Constantino et al., 2021). The FRN presents more negative amplitudes for negative (e.g., loss) feedbacks when compared with positive feedbacks (e.g., win) (Gehring & Willoughby, 2002; Glazer et al., 2018; Nieuwenhuis et al., 2005; Sambrook & Goslin, 2015), is considered an early index of reward outcome evaluation, and seems to be modulated by the affective impact of an event (e.g., Boksem & De Cremer, 2010). The P3b component is a positive deflection captured in the centroparietal electrodes in between 300 to 600 ms post feedback presentation (e.g., Constantino et al., 2021). The P3b is modulated by reward magnitude, valence, and probability of an outcome (Hajcak et al., 2005; Mei et al., 2018). The P3b is thought to index information integration and categorization (Kok, 2001; Pisauro et al., 2017; Wichary et al., 2017), and deliberate/active working memory dynamics (Chase et al., 2011; Eppinger et al., 2017; Polich, 2007) responsible for communication within the brain’s frontal networks involved in decision-making (Kok, 2001; Nieuwenhuis et al., 2005; Twomey et al., 2015; Warren & Holroyd, 2012). Both FRN and P3b components have been useful for the understanding of outcome evaluation in economic decision-making settings and therefore show promise for a better understanding of attack and defence conflict in an economic predator-prey contest such as the Predator-Prey game.

### General Goal and Hypotheses

The main goal of the present ERP study is to characterize the neurophysiological correlates of feedback processing (FRN and P3b) in a decision-making task designed to operationalize attack and defence conflicts over resources, and thus providing a better understanding of social interactions under conflict. Given previous data on the behavioural and psychological determinants of attack and defence we hypothesize the following: (H1) the behavioural investment in attack will be higher, and slower than in defence; (H2) participants will show increased FRN and P3b amplitudes to losses when compared wins in both attack and defence; (H3) FRN amplitudes will be higher for defence when compared with attack; and (H4) P3b amplitudes will be higher for attack when compared with defence.

## 2. Methods

### 2.1. Participants

Fifty community-dwelling participants without history of neurological disease were recruited through social media and through faculty’s official website. Nevertheless, only 47 volunteers (28 female) were considered for the further statistical analysis of the study due to dropouts and non-comprehension of the tasks involved in the experimental protocol. Participant’s age ranged between 19-45 years old (*M* = 24.36, *SD* = 5.24) and years of education ranged between 12-21 years (*M* = 15.57, *SD* = 2.08).

From the 47 participants included in the behavioural data analysis, ERP data from 11 participants playing the attacker role and ERP data from 18 participants playing the defender role were excluded due to excessive background and movement noise in the EEG data.

### 2.2. Materials

#### Predator-Prey Game

The Predator-Prey Game (PPG) is an economic decision-making contest adapted from De Dreu and colleagues (2015). The contest operationalizes conflict in the form of attack and defence where players are assigned to one of the two roles. On each trial, the task started with a fixation point with 700 ms duration followed by a decision slide where participants should decide how many credits to invest, ranging from 0 to 9 out of an initial endowment of 10 credits. After participants made their decision, a fixation point was presented for a random time-interval between 1000 and 2000 ms followed by the feedback on that interaction, exhibited for 5000 ms, followed by a 2000 ms slide displaying the quantity of credits won in that trial. The result of the interaction was computed on a trial-by-trial basis comparing the response of the participant with a random opponent’s response, whose probability was determined by the weights of possible responses given by the Nash Equilibrium (for a more detailed description of this procedure, please see De Dreu et al., 2015). The trial structure is displayed on Figure 1. Participants played 50 trials as attackers, and 50 trials as defenders in two separate sessions. As behavioural output of the task, the mean investment in 50 trials for each role and the mean decision time used to make the behavioural decision were computed.

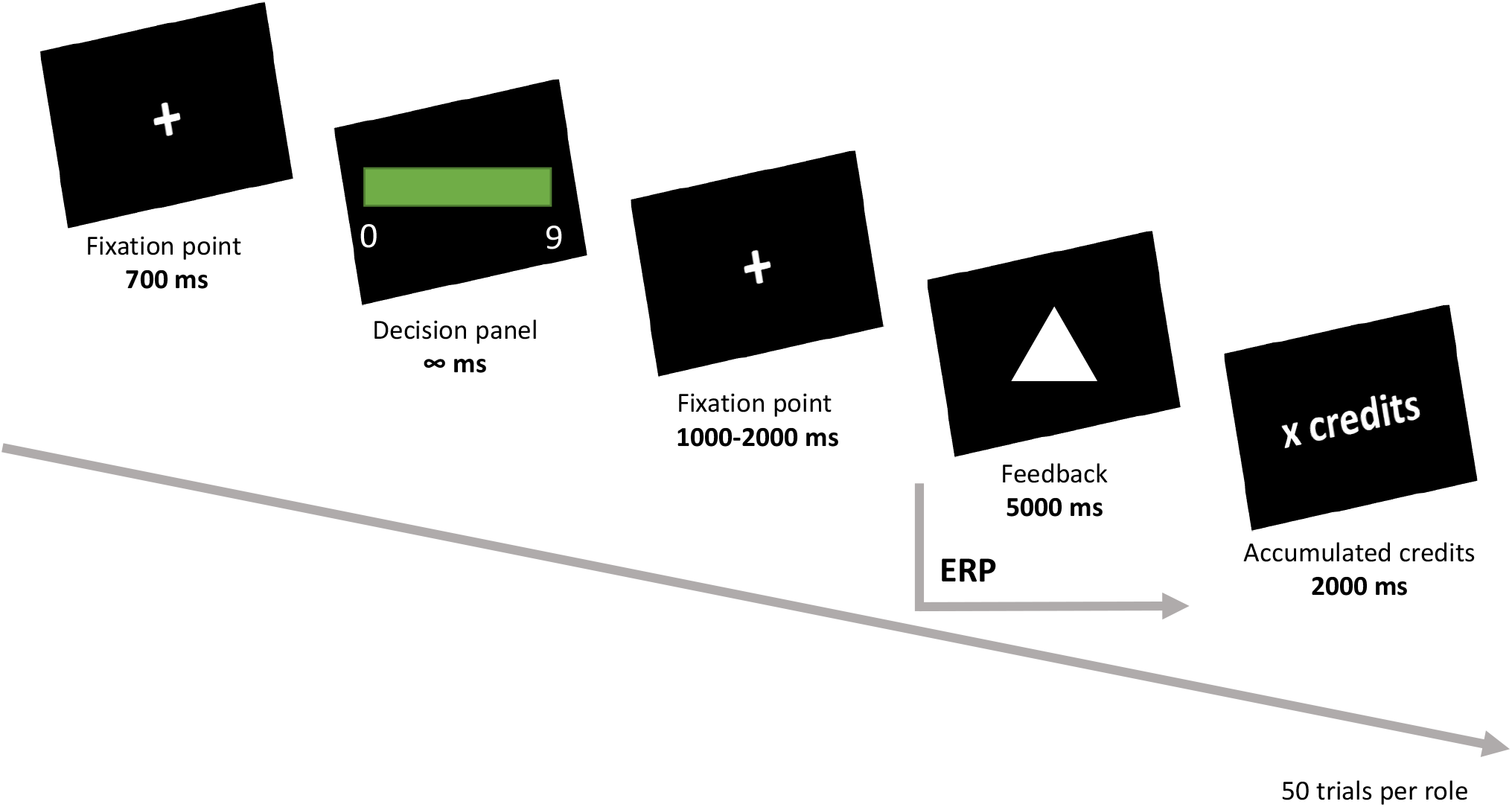

The outcome of each interaction was determined differently based on the role participants were playing. For attackers, in case the investment in predation was bigger than the opponent’s investment in defence, the attacker would accumulate his non-invested credits plus the non-invested credits of the defender. When playing as defender, whenever the investment in defence was bigger or as high as the opponent’s investment, the defender would accumulate the remaining non-invested credits. For attackers, a loss feedback was displayed whenever the investment was lower than the opponents or whenever the accumulated credits were lower than the initial endowment of 10 credits. For defenders, a loss feedback was displayed whenever participants invested less than the opponent. To prevent possible priming effects potentiated by the ‘attacker/defender’ wording, when assigning the role in which participants were playing, the instructions only reflected the mathematical rules of the interaction (De Dreu & Kret, 2016), As so, participants were fully informed on the mathematical rules underlying each interaction and a set of multiple choice questions were formulated to ensure comprehension of the dynamics of each role.

### 2.3. EEG data collection, signal processing and ERP measures extraction

The electroencephalographic data was recorded using a high impedance NetAmps 300 system from Electrical Geodesics Inc. (USA), with a 128-channel Hydrocel Geodesic Sensor Net. The signal was digitized at 500 Hz with a vertex reference (Cz), and the electrode impedances were kept below 50 KΩ. During acquisition, an antialiasing filter was automatically applied by the acquisition software (Net Station v4.5). The filter was a Butterworth low-pass, designed to have a frequency response as flat as mathematically possible in the passband, rolling off toward zero in the stopband at 250Hz (the Nyquist frequency of the selected sampling rate). Data processing and analysis was conducted using EEGLAB toolbox v13.6.5b (Delorme & Makeig, 2004) and the ERPLAB plugin v6.1.3 (Lopez-Calderon & Luck, 2014). EEG signals were then resampled to 250 Hz and bandpass filtered, using a causal bandpass filter with 0.1 Hz as the low half-amplitude cutoff (transition band width of 0.1 Hz and roll-off at 12 dB/octave) and 30 Hz as the high-amplitude cutoff (transition band width of Hz and roll-off at 12 dB/octave). After filtering, channels with excessive noise were removed, never exceeding a number of channels > 10% of the total number of electrodes. The EEG was then submitted to a temporal independent component analysis (ICA, Jung et al., 1998). Independent components (ICs) identified as corresponding to eye blinks, saccades, or heart rate were removed, in order to correct for this type of artifacts. The artifact correction procedure was the following: (a) visual inspection of the ICs and pre-selection of those resembling to correspond to eye blinks, saccades, and heart rate activity; (b) visual inspection of the time course of the pre-selected ICs; (c) comparison of the original signal with the back-projected signal without the selected components; (d) if the changes in the corrected signal were circumscribed to the latency of the specific artifacts without changing the EEG morphology at other latencies, the components were removed; if not, no correction was performed at this stage (i.e., remaining artifacts were later removed during visual inspection). After artifact correction, deleted channels were interpolated by means of spherical spline interpolation, which weights each electrode spatially in a way that is qualitatively consistent with dipole fields (Ferree, 2006). The signal was then re-referenced to the average of all electrodes. Finally, the EEG signal was segmented into epochs around the onset of the stimulus of interest: 1000 ms epochs with 200 ms before (baseline) and 800 ms after the onset of the feedback slide. All segments were subjected to visual inspection, and epochs containing artifacts not corrected using ICA were deleted. The epoch deletion decision was based on visual inspection (i.e., based on EEG morphology) and performed by consensus of two experts. The remaining epochs were baseline corrected (200 ms baseline duration) and averaged into the conditions of interest (i.e., Gain and Loss feedback). The time windows for FRN and P3b extraction were defined by visual inspection of the morphology of the ERP and are within the time intervals defined in the literature for both FRN and P300 components (Hajcak et al., 2007). The following ERP component measures were computed: (1) FRN amplitude, defined as the mean amplitude of the 50 ms time window around the most negative peak in the 80 to 250 ms time-window post feedback presentation at FCz; and (2) P3b amplitude, defined as the mean amplitude of the 50 ms time-window around the most positive peak in the 300 to 450 ms time-window post feedback presentation at Cz and Pz electrodes. As we used a high-density EEG setup, channels of interest were defined as the regional average of locations defined by the extended international system 10-5: FCz as the average of the electrodes E5, E6, E7, E12, E13, E106, and E112; Cz as the average of the electrodes E7, E31, E55, E80, E106, and E129; and Pz as the average of the electrodes E61, E62, E67, E72, E77, and E78.

### 2.4. Procedure

Participants were invited to engage in a two-session data collection, apart from each other between five to ten days – one session to play as attacker and other as defender (the oreder was counterbalanced between participants). On each session, upon arrival at laboratory, participants were informed that the main goals of the study was to analyse how people make decisions and gave the written informed consent to engage in the study. The EEG signal was recorded while participants were seated in a chair one meter away from a 17’’ computer monitor, where the computerized version of the PPG was delivered with the E-Prime 2.0 software (Psychology Software Tools, Pittsburgh, PA). At the end participants receive a monetary compensation for the time spent in the experiment. The task and procedures were approved by the local ethics committee, and all participants were treated in accordance with the 2013 revision of the declaration of Helsinki.

### 2.5. Statistical Analyses

The statistical analyses were performed using IBM SPSS Statistics 26 (IBM Corporation, Armonk, NY, USA). Descriptive statistics for demographic, behavioural, and physiological variables are presented. The behavioural and neurophysiological correlates of the Predator-Prey Game were assessed using Repeated Measures ANOVA. Whenever an outlier was identified from standardized residuals (> 2.0 or < -2.0) of the ANOVA model, participants were excluded (Everitt, 1998). Multiple comparisons were performed using the Bonferroni correction for multiple comparisons and the alpha threshold considered for all analyses was .05.

## 3. Results

The descriptive statistics for the behavioural and ERP data are described in Table 1.

**Table 1.** Descriptive statistics of the behavioural (investment and response time) and ERP correlates (FRN and P3b peak amplitudes) of the PPG.

|  | Attack |  |  | Defence |  |  |
| --- | --- | --- | --- | --- | --- | --- |
|  | N | Mean (SD) | Min - Max | N | Mean (SD) | Min - Max |
| Investment | 43 | 4.79 (1.07) | 2.18 – 6.88 | 43 | 5.00 (0.91) | 3.54 – 7.38 |
| Response Time (s) | 41 | 1.78 (0.8) | 0.39 – 3.81 | 41 | 1.24 (0.56) | 0.52 – 2.91 |
| FRN Gains (FCz) | 21 | -2.08 (1.42) | -5.87 – 0.00 | 21 | -2.21 (1.20) | -3.99 – 0.04 |
| FRN Losses (FCz) | 21 | -2.46 (1.42) | -5.53 – 0.08 | 21 | -2.96 (1.15) | -5.19 – -0.80 |
| P3b Gains (Cz) | 16 | 4.56 (2.07) | 1.37 – 8.96 | 16 | 2.29 (1.67) | -0.20 – 5.28 |
| P3b Losses (Cz) | 16 | 4.81 (2.58) | -0.28 – 8.94 | 16 | 3.56 (2.54) | -0.75 – 8.78 |
| P3b Gains (Pz) | 16 | 4.91 (2.09) | 0.70 – 8.72 | 16 | 2.96 (1.61) | 0.79 – 7.42 |
| P3b Losses (Pz) | 16 | 4.80 (1.56) | 2.87 – 7.55 | 16 | 2.37 (2.18) | -2.33 – 6.69 |

*Note.* SD: standard deviation; Min: minimum; Max: maximum.

### 3.1. Behavioural correlates of the PPG

Repeated Measures ANOVA with *Role* (Attack, Defence) as within-subjects factor and *Order* (Attack-Defence, Defence-Attack) as a between subjects factor for mean investment revealed the following: a non-significant effect of *Role* [*F*_(1,41)_ = 1.19; *p* = .281], and a non-significant *Role*Order* interaction [*F*_(1,41)_ = 4.03; *p* = .051]. Nonetheless, Bonferroni multiple comparisons test on the *Role*Order* interaction revealed participants that played as defenders first (defence-attack) tend to invest more in defence (*M* = 4.98; *SEM* = 0.21) than in attack (*M* = 4.31; *SEM* = 0.22; *p =* .040). Participants that played as attackers first (attack-defence) made similar investments in attack (*M* = 5.21; *SEM* = 0.21) and defence (*M* = 5.01; *SEM* = 0.19, *p =* .506). For response time, the same analysis revealed a significant effect of *Role* [*F*_(1,39)_ = 17.3; *p* < .001; *η*^*2*^_*p*_= .31] with participants showing faster response times for defence (*M =* 1.26; *SEM* = 0.08) when compared with attack (*M =* 1.76; *SEM* = 0.12), and a significant *Role*Order* interaction [*F*_(1,39)_ = 23.2; *p* < .001; *η*^*2*^_*p*_ = .37]. Bonferroni multiple comparisons test on the *Role*Order* interaction revealed participants that played as attackers first are faster to invest in defence (*M* = 0.98; *SEM* = 0.11) than in attack (*M* = 2.07; *SEM* = 0.16; *p <* .001). Participants that played as defenders first showed similar response times in attack (*M* = 1.45; *SEM* = 0.17) and defence (*M* = 1.53; *SEM* = 0.11, *p =* .656). See Figure 2 for a visual depiction of the comparisons regarding mean investment and response time in attack and defence.

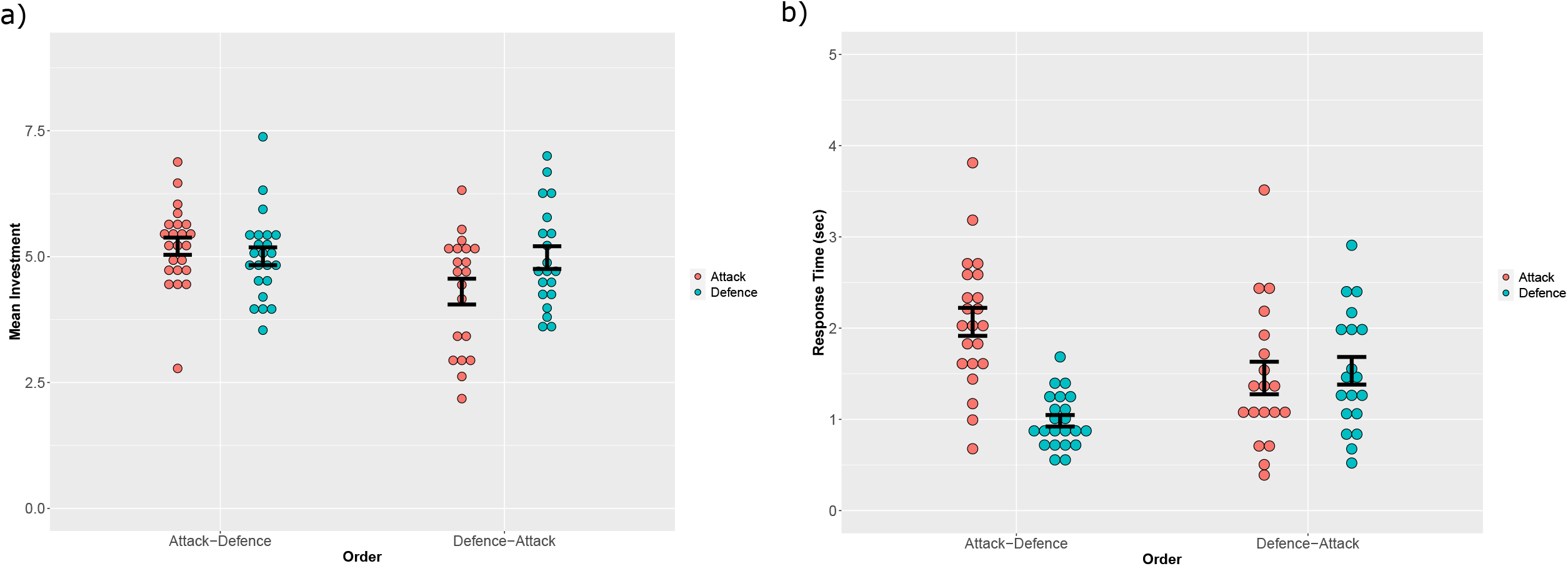

### 3.2. Electrophysiological correlates of the PPG

The ERPs elicited by the feedback at FCz and Cz are depicted in Figure 3.

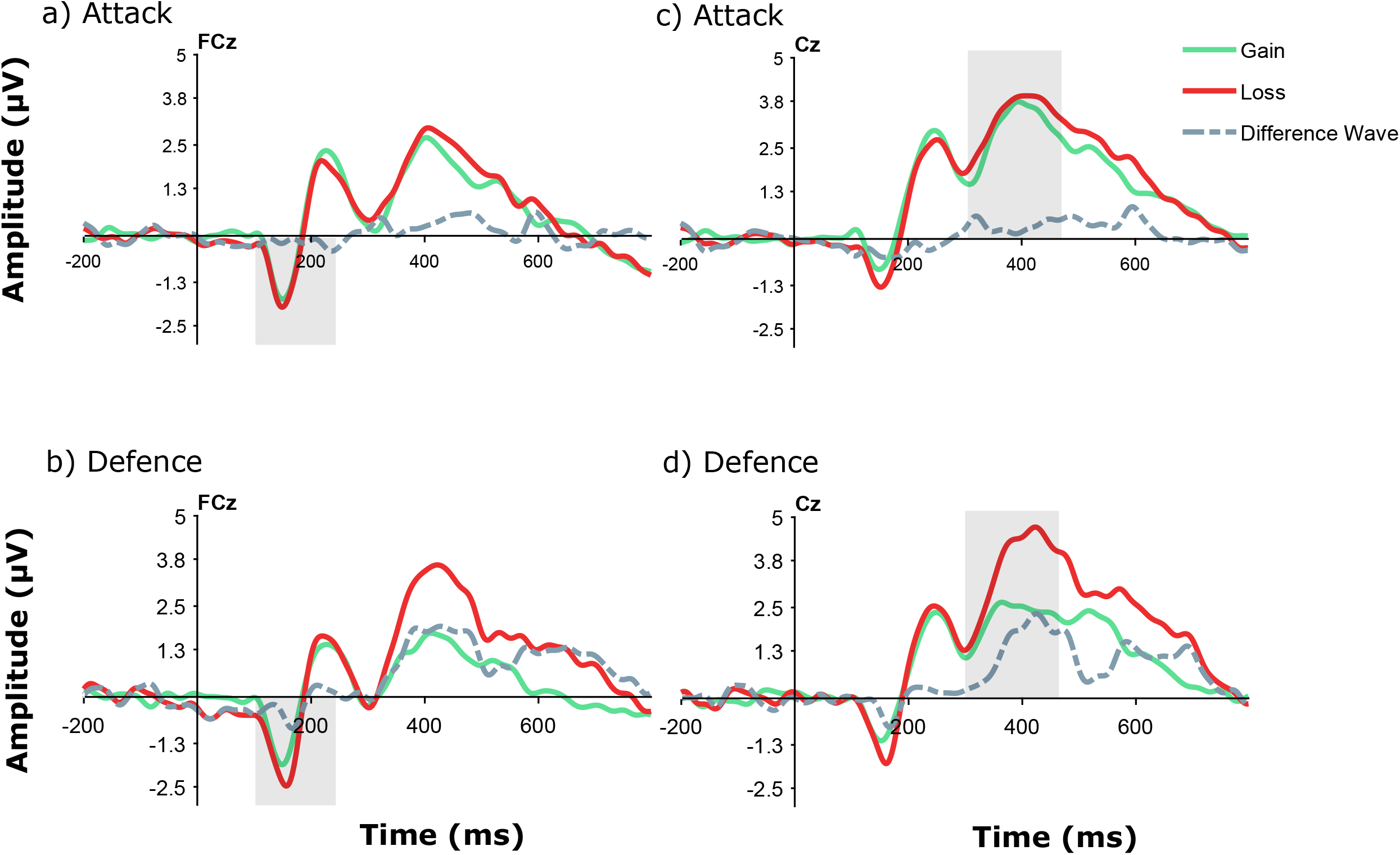

For the FRN, a Repeated Measures ANOVA with *Role* (Attack, Defence) and *Outcome* (Gain, Loss) as within-subjects factor, and *Order* (Attack-Defence, Defence-Attack) as a between subjects factor revealed a significant main effect of *Outcome* [*F*_(1,19)_ = 6.60; *p* = .02; *η*^*2*^_*p*_ = .25] with losses eliciting increased FRN amplitudes (*M* = -2.76; *SEM* = 0.20) than gains (*M* = -2.19; *SEM* = 0.22). No main effect of *Role* [*F*_(1,19)_ = 2.17; *p* = .16] nor *Role*Outcome* [*F*_(1,19)_ = 0.72; *p* = .41], *Role*Order* [*F*_(1,19)_ = 2.38; *p* = .14], *Outcome*Order* [*F*_(1,19)_ = 0.06; *p* = .81], nor *Role*Outcome*Order* [*F*_(1,19)_ = 1.90; *p* = .18] interactions were found.

For P3b, a Repeated Measures ANOVA with *Role* (Attack, Defence), *Outcome* (Gain, Loss), and *Channel* (Cz, Pz) as within-subjects factor, and *Order* (Attack-Defence, Defence-Attack) as a between subjects’ factor revealed a main effect of *Role* [*F*_(1,14)_ = 70.2; *p* < .001; *η*^*2*^_*p*_ = .83] with higher P3b amplitudes for attack (*M* = 4.77; *SEM* = 0.29) when compared with defence (*M* = 2.79; *SEM* = 0.37), and a significant *Channel*Outcome* interaction [*F*_(1,14)_ = 6.61; *p* = .022; *η*^*2*^_*p*_ = .32]. However, Bonferroni multiple comparisons test did not reveal any differences in the P3b amplitudes for gain and losses at Cz and Pz channels. No main effect of *Outcome* [*F*_(1,14)_ = 0.38; *p* = .55], *Channel* [*F*_(1,14)_ = 0.14; *p* = .91], nor *Channel*Role* [*F*_(1,14)_ = 0.70; *p* = .42], *Role*Outcome* [*F*_(1,14)_ = 0.24; *p* = .63], nor *Role*Outcome*Channel* [*F*_(1,14)_ = 2.44; *p* = .14] interactions were found. The *Order* factor (between-subjects) did nor interact the within-subjects factors nor with the 2^nd^ and 3^rd^ order interactions (all *p >* .05).

## 4. Discussion

Interpersonal and social conflict is present in distinct levels of human society and has the potential to affect, as an example, nations welfare and individual life trajectories (De Dreu & Gross, 2019). The study of the brain correlates of conflict has the potential to unveil the mechanisms underlying conflict. At this level, game theory approaches are fundamental tools to operationalize the subjective value in conflict interactions and the asymmetric operationalization of the Predator-Prey contest provides an operationalization that resembles a subset of interpersonal conflicts present social groups were individuals compete with distinct motivations (e.g., profit vs loss avoidance; De Dreu et al., 2015, 2019; De Dreu & Gross, 2019) The present study aimed at characterizing the behavioural and brain ERP correlates of interpersonal conflict over resources under an attack-defence economic decision-making formulation of the Predator-Prey Game. Specifically, we focused on the FRN and P3b components of the ERP responses to outcome feedback in attack-defence interactions.

The strategies and brain networks differently recruited for attack and defence support the notion that distinct mechanisms underlie attack and defence: while defence seems to be more automatic in nature, eliciting increased activation of limbic and anterior brain regions responsible for threat detection, attack relates with increased recruitment of prefrontal brain regions linked with top-down control and deliberation (De Dreu & Gross, 2019). As hypothesized, our behavioural results partially support the notion that defence is motivationally more relevant and more automatic than attack, with individuals showing increased investment and shorter response times for defence when compared with attack, which is in line with previous reports on the PPG (De Dreu et al., 2015, 2019, 2021; De Dreu & Gross, 2019; De Dreu & Kret, 2016; van Dijk & De Dreu, 2021). Importantly, this behavioural pattern only emerged when controlling for the order of play, meaning that individuals invested more in defence when playing as defenders first and showed reduced response times for defence when playing as attackers first. Our repeated measures design included two sessions with at least one week of interval between them, where participants played one role in day 1 (e.g., attack) and the other in day 2 (e.g., defence), with the order being counterbalanced between participants. Previous studies using the PPG typically employed a similar repeated measures design with the two roles being playing at the same day (e.g., De Dreu et al., 2015). Our results suggest that even a minimum of one week interval between sessions is not sufficient to reduce the recency effect typical of behavioural decision-making task with repeated measures designs in distinct contexts (e.g., Highhouse & Gallo, 1997; Nofsinger & Varma, 2013), which should be considered for future experimental implementations of the PPG.

At the level of the FRN and P3b correlates of feedback processing in the PPG, our results partially support our hypothesis. While FRN showed increased amplitudes for losses when compared with gains, the FRN modulations by attack and defence were similar. On the other hand, P3b showed similar modulations for losses and gains, with increased P3b amplitudes for attack when compared with defence. As so, while the early (and more automatic) reactivity to losses was present in both attack and defence, the later allocation of attentional resources indexed by the P3b was more pronounced for attack, thus partially supporting the notion that attack is more cognitively demanding, and thus more deliberate, than defence (De Dreu & Gross, 2019). Previous studies suggest that the FRN seems to indicate a binary evaluation of “*good*” vs “*bad*” outcomes (e.g., Hajcak et al., 2006, 2007), and our results suggest that this early evaluation presents similar biological relevance for attack and defence. It is import to note however, that although within the range of the time windows previously used to extract the FRN, the frontocentral negativity differentiating losses and gains emerged earlier than in previous studies using the FRN (e.g., Hajcak et al., 2005, 2006, 2007), and future replications of our findings are necessary to assess the robustness of the associations reported here.

## Conclusion and Future Directions

The present study provides for the first time a characterization of the ERP correlates underlying feedback processing in attack and defence social interaction conflict. The integration of the behavioural and ERP findings support the theoretical model positing that attack elicits more deliberate and less automatic processes than defence (De Dreu et al., 2019, 2021). Notwithstanding the significance of the reported findings future studies should test the robustness of the reported finding in future replications. Abnormal social interaction patterns are characteristic of several psychopathological and personality manifestations, such as anxiety, social phobia, psychopathy and antisocial behaviour (Jones et al., 2007; Plana et al., 2014; Sharp & Vanwoerden, 2014). Game theoretical approaches to social interactions suggest a framework for the analysis of composite patterns of behaviour in the form of interpersonal interactions (Camerer, 2003), where asymmetrical interactions, such as the Predator-Prey conflict provide an operationalization similar to several human interaction conflicts (De Dreu & Gross, 2019). Thus, a better understanding of the brain mechanisms underlying not only feedback processing, but also subjective value computations in attack and defence conflict offers a promising venue for a better understanding of maladaptive social interaction patterns.

